# Mechanical Hand Synergies during Dynamic Hand Movements are Mostly Controlled in a Non-Synergistic Way by Spinal Motor Neurons

**DOI:** 10.1101/2023.07.25.550369

**Authors:** Marius Oßwald, Andre L. Cakici, Daniela Souza de Oliveira, Dominik I. Braun, Alessandro Del Vecchio

## Abstract

Precise control of spinal motor neurons is crucial for voluntary hand and digit movements. However, the specific mechanisms by which motor unit ensembles govern dynamic synergistic and individual digit tasks remain poorly understood. We recorded synchronized 3D hand kinematics and high-density surface EMG (HD-sEMG) data from extrinsic hand muscles of twelve participants during 13 dynamic hand and digit movement tasks, consisting of single-digit flexion/extension and mechanically synergistic grasping tasks.

We extracted single motor unit (MU) activity and identified identical MUs across tasks. We extracted 7.8 ± 1.8 MUs per task and participant and found 182 out of 554 total MUs active during multiple movements. Analysis of the MU discharge patterns revealed two groups of motor units that were categorized into *prime mover MUs*, showing strong correlation between firing rate modulation and digit kinematics, and *postural MUs* with little modulated activity. We found these motor units could switch between the two modes, showing either postural or movement encoding activation depending on the task. However, MUs acted as prime mover only for one specific digit. We further observed highly task specific recruitment of *prime mover* MUs. Across participants, we found only 9 ± 8.2 % of *prime mover* MUs active during a grasp task and any single digit task involved in the grasp motion.

We draw three conclusions: (1) Single digits are controlled by distinct groups of MUs. (2) Unexpectedly, mechanically synergistic grasp movements are mostly controlled in a non-synergistic way by distinct groups of MUs. 3) Multiple manifolds construct the movement of the human hand, and each motor unit can flexibly switch between postural and dynamic modes.

**Significance Statement:** We investigated the neural control of motor unit ensembles during single-digit and synergistic grasping tasks in dynamic conditions. We found that motor units exhibited strong movement-correlated activity only for one specific digit. We further observed highly task specific recruitment of motor units during mechanically synergistic grasp movements, showing that on a motor unit level, mechanically synergistic movements are controlled in a non-synergistic way. The findings extend the knowledge of motor unit recruitment strategies in natural movements and have strong implications in the field of neurorehabilitation and control of assistive devices.

## Introduction

The human hand is a highly complex apparatus both on the mechanical and neuromuscular level with a relatively high-dimensional functional output. Although such a high variety of postures and movements in a high number of degrees of freedom can be performed, kinematic recordings of the hand posture reveal a large portion of the postural variance can be explained by only few latent embeddings (Della Santina et al., 2017; Grinyagin et al., 2005; Mason et al., 2001; Santello et al., 1998). However, it is unknown if the dimensionality of the output of spinal motor neurons is reduced in a similar extent as the kinematical output. It may be assumed that the components accounting for just little amounts of the functional variance, are explained and controlled by specific motor neurons encoding distinct tasks. How rigid or flexible these motor neurons are controlled is still in discussion.

Marshall et al. (2022) observed flexible control of motor units (MUs) while varying mechanical conditions in isometric contractions, suggesting a more flexible control of motor neurons than previously documented (Henneman, 1957). Most research targeting the synergistic control of muscles and MUs during varying mechanical conditions, hand postures and synergistic grasping tasks in humans or primates is performed in isometric conditions (Bräcklein et al., 2022; Del Vecchio et al., 2022; Feeney et al., 2018; Marshall et al., 2022; Ricotta et al., 2023; Tanzarella et al., 2021; Vecchio et al., 2023). Extending the investigations to dynamic movements adds nonlinear components to the recruitment and firing behavior of motor units, that may be further changed with the speed of contraction (Theeuwen et al., 1994). The lengthening and shortening of muscles in dynamic conditions generates afferent inputs that strongly modulate the neural input received by the MUs. There are no studies investigating the dimensionality of control for a representative population of motor units during unconstrained hand and digit movements at different movement speeds across the full degrees of freedoms of the human hand.

We developed a framework for camera-based, markerless acquisition of 3D hand kinematics, covering all degrees of freedom of the hand with precisely synchronized high-density surface electromyography (HD-sEMG) data from the extrinsic hand muscles during a series of hand and digit movements. We decoded the activity of spinal motor neurons during isolated single digit movements at different speeds and mechanically synergistic grasping motions with the goal to understand the recruitment strategies and activation patterns of identical MUs across movement tasks. We found that all movements of the human hands are mostly controlled by distinct populations of motor neurons. Single digits are controlled by distinct groups of motor units. Unexpectedly, mechanically synergistic grasp movements are primarily governed by separate groups of motor units, lacking a synergistic pattern at the motor unit level.

The presented research is valuable for understanding the correspondence of motor unit activity and movement, possibly leading to advances in the field of neurorehabilitation through better detection of movement intentions for the control of assistive devices since most of neural lesions still preserve spared functional modulated motor unit activity (Oliveira et al., 2022; Ting et al., 2021).

## Results

### Framework for synchronized markerless 3D kinematics acquisition and motor unit data

We developed a framework for markerless acquisition of 3D hand kinematics with synchronized recording of HD-sEMG data (Fig. 1A). The participants hand movements were recorded with multiple industrial laboratory RGB cameras. The hand positions were extracted by a custom neural network trained to track 21 labels on hand and digit joints in all videos to obtain the respective 2D kinematics. We then triangulated the 2D positions to get the 3D positions of the 21 labels in all video frames. Figure 1 A shows the schematic workflow of the 3D kinematics and HD-sEMG acquisition. We recorded data from 12 participants performing 13 dynamic hand and digit tasks, consisting of single-digit flexion and extension of all five digits at low (0.5 Hz) and high (1.5 Hz) movement frequency (and therefore movement speed) and three grasping tasks (two-digit precision pinch using thumb and index digit, three-digit precision pinch using thumb, index and middle digit, five-digit power grasp). The grasping tasks were chosen to combine the above-described single-digit tasks in their same degrees of freedom (flexion/extension). Video S2 shows exemplary experimental videos with the tracked and triangulated 3D kinematics.

**Figure 1.**
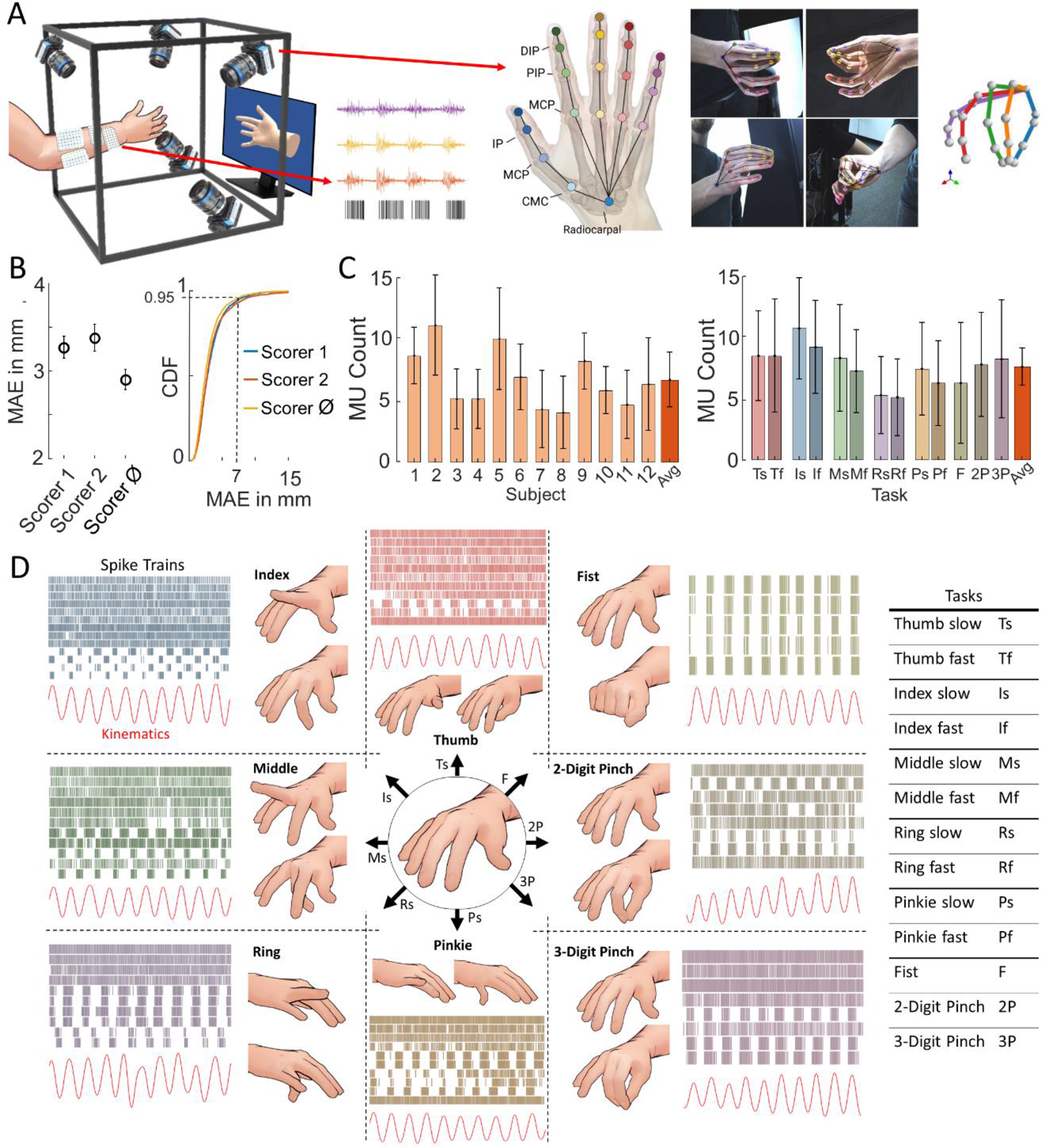
Synchronised HD-sEMG/motor unit data and 3D kinematics acquisition. (A) Cameras recorded video from different angles. A neural network tracked 21 labels, corresponding to hand and digit joints and fingertips, in all videos. Subsequent triangulation results in the 3D kinematic data. (B) Left: Mean absolute error (MAE) between 3D positions obtained from neural network and manually labelled videos by two experts. Right: Cumulative distribution function of the MAE showing 95% probability for a 3D MAE of any label of < 7 mm. (C) Average number of found motor units per participant (left) and task (right). (D) Motor unit firings and digit tip kinematics of one participant of single-digit periodic flexion/extension tasks at 0.5 Hz and grasp tasks and list of all performed task and their hereafter used abbreviation.

The HD-sEMG data was decomposed into single motor unit activity with a gradient convolution kernel compensation algorithm (Holobar & Zazula, 2007). Only motor units with a pulse-to-noise ratio (PNR) of >30 dB and a minimum of 100 detected action potential firings per task were included in the further analysis. We then checked the reliability in tracking the same action potential waveform across tasks with multiple analyses (see Fig. 2 and Fig. S1). We found the motor unit action potential (MUAP) shapes and location of MUs to be consistent throughout the tasks, with distinct hotspots of activation for each motor unit (Fig 2., Fig. S1).

**Figure 2.**
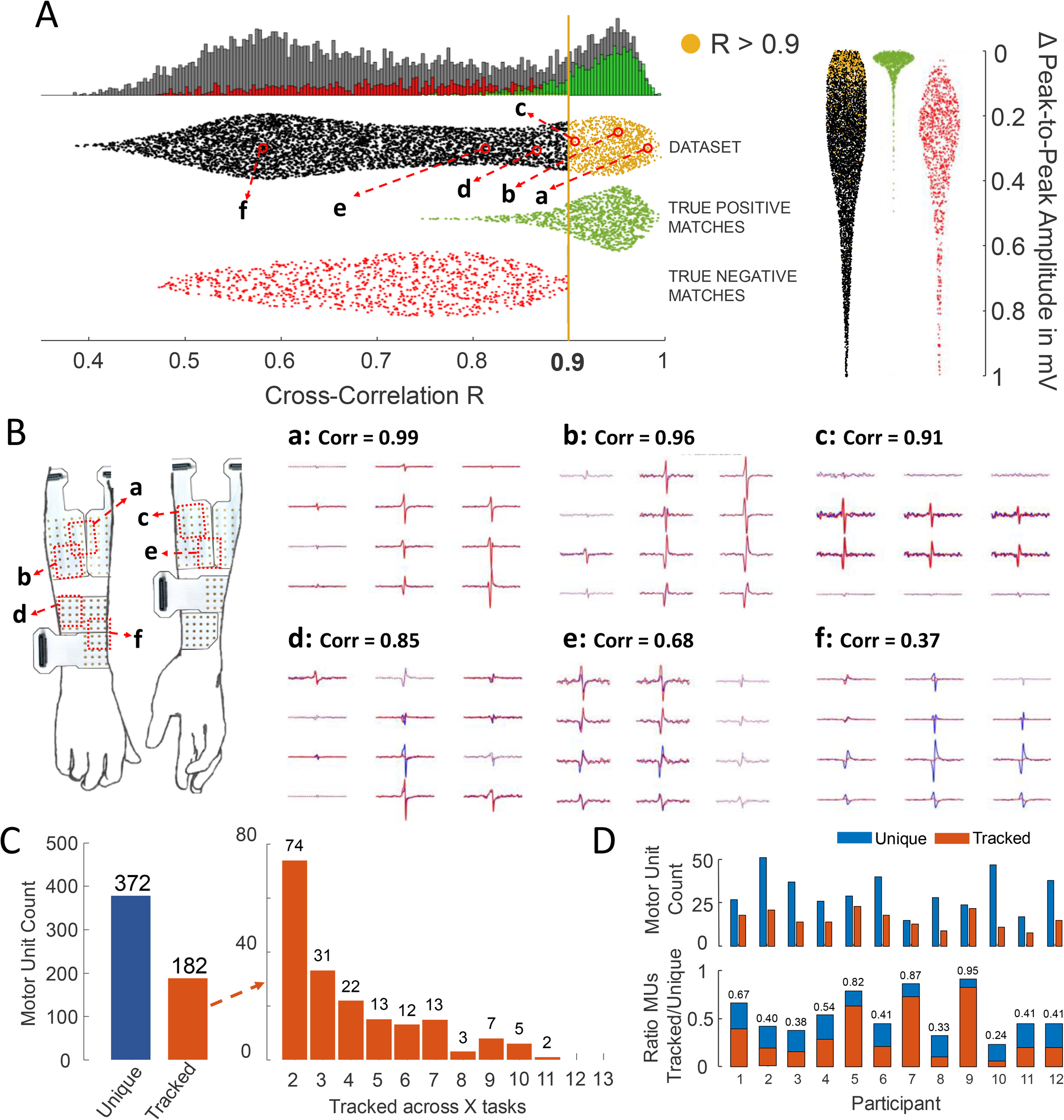
Tracking of motor units across dynamic movement tasks. (A) Left: Distribution of cross-correlation values R of MUAP shapes of pairs of MUs obtained from all possible combinations of two MUs of the same participant in the dataset (black), of MUAP shapes taken from the first and second half of the signal of all MUs as a known group of identical Mus (green, True positive matches) and of pairs of MUs from the same task as a known group of distinct MUs (red, true negative matches). Chosen decision boundary of R=0.9 for positive matches of MU pairs marked in yellow. Right: Distribution of difference in peak-to-peak amplitudes of MU pairs in mV. MU pairs with R values > 0.9 marked in yellow. (B) MUAP shapes of a subset of the full HD-sEMG grid of six exemplary MU pairs taken from the full range of R values for demonstration of MUAP similarity. (C) Left: Number of MUs found in only one task (Unique MUs) and MUs tracked across a minimum of two tasks (Tracked MUs). Right: Breakdown of tracked MUs into the number of tasks each MU could be identified in. (D) Top: Number of MUs found for each participant, split into unique (blue) and tracked (red) MUs. Bottom: Ratio of tracked vs. unique MUs for each participant.

We found an average of 7.8 ± 1.6 motor units per task and 7.8 ± 2.5 motor units per participant with an average PNR of 34.9 ± 1.6 over all participants and tasks (Fig. 1D and Table 1). The mean firing rate (MFR, pulses per second, pps) of motor units was 12.5 ± 1.6 pps across participants and 12.5 ± 0.7 pps across tasks with no significant difference in MFR between fast and slow movement tasks (Table 1).

**Table 1.**
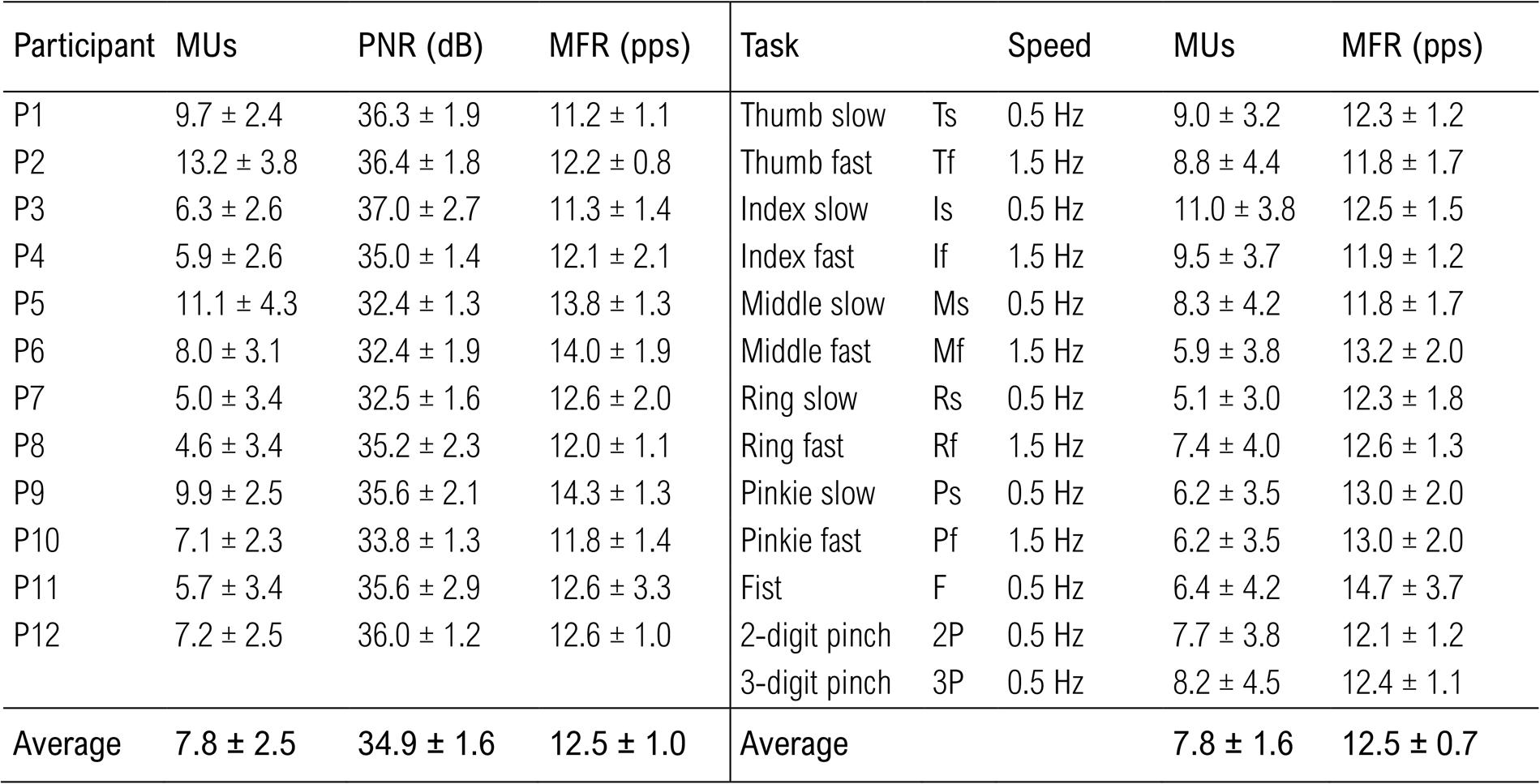
Motor Unit Decomposition. Left: Number of identified motor units, pulse-to-noise ratio (PNR) and mean firing rate (MFR) across participants, averaged over tasks. Right: Number of identified motor units and MFR across tasks, averaged over participants.

Figure 1E displays the performed movements, exemplary spike trains, and digit tip kinematics of one participant for the single-digit tasks performed at 0.5 Hz frequency and grasping movements. In all further figures and analyses, a positive slope of the kinematics corresponds to digit extension with the local maximum at the peak extension of that movement cycle. A negative slope corresponds to digit flexion with the local minimum at the peak flexion of that movement cycle. In grasping tasks, the digit tip kinematics of all involved digits were combined into one representative movement trajectory with positive and negative slopes corresponding to extension and flexion, analogue to the single-digit tasks.

#### Motor unit tracking across tasks

Motor units were tracked across tasks to identify the activity of identical motor units during different tasks. The tracking was based on comparing the motor unit action potential shapes of MUs found in different tasks. Cross-correlation R was computed between the MUAPs of all possible combinations of two MUs within the identified population of MUs of a participant. Only MU pairs with a cross-correlation value R of >0.9 were considered as positive matches. This value was chosen by computing the distribution of R values within known groups of identical (true positive matches) and distinct MUs (true negative matches, see Fig. 2A). This value is considerably higher than previously reported in the literature for longitudinal tracking of MUs in isometric conditions (Martinez-Valdes et al., 2017; Vecchio & Farina, 2019), due to the fact that we employed a larger number of electrodes and improved the tracking by extracting the correlations from the motor unit specific territory (see Methods section for further details). This analysis ensured no false positive matches in the resulting pool of tracked MUs. We further validated the tracking by looking at differences in the peak-to-peak amplitudes (ΔP2P) of the motor units (Fig. 2A). We consistently found a strong clustering of R and ΔP2P values, so that there was little difference in the amplitude values of the motor unit action potential waveforms across the tracked MUs (Fig. 1A). Figure 2B shows comparisons of MUAP shapes of pairs of MUs across the full range of observed cross-correlation values. Out of all 554 motor units identified across all participants and tasks, 182 MUs could be tracked across at least two tasks (from now on considered as *tracked MUs*). The remaining MUs (n=372) could not be tracked and were therefore found active only in a single task (*unique MUs*). Figure 2C shows the distribution of the number of tasks the tracked MUs were identified in. The majority of tracked MUs were identified across few tasks. However, some motor units showed activity during a large range of movements, with a maximum number of 11 tasks (n=2). The ratio of tracked/unique MUs split into participants is shown in Figure 2D (Tracked/Unique MUs= 0.53 ± 0.23).

### Individual digit flexion and extension movements do not share *prime mover* motor units

We used the coefficient of variation (CoV) as a measure of task modulation of motor unit firing activity (Fig. 3A). We calculated the CoV on the smoothed firings and report details on the smoothing method. Figure 3B shows a matrix of all motor units of one participant that have been identified in at least two tasks. The color indicates the task modulation of the firing behavior for each task according to the respective CoV value. Figure 3C displays a subset of the matrix with the individual CoV values. Motor units with a CoV of >0.5 in at least one of their identified tasks are hereafter referred to as “prime mover” MUs, as their firing pattern is strongly correlated to the digit tip trajectory (R=0.91 ± 0.063 across all participants and tasks) with high activity during either the flexion or extension period of the movement and only weakly or not active during the respective other period. Therefore, the prime mover motor units are units that encode the movement of the digit. There was also a large proportion of motor units that did not correlate with any of the task kinematics, resulting in CoV values of <0.5 (Fig. 3C). These units are hereafter referred to as “postural motor units” and encode the static posture of the hand with virtually no discharge rate modulated activity.

**Figure 3.**
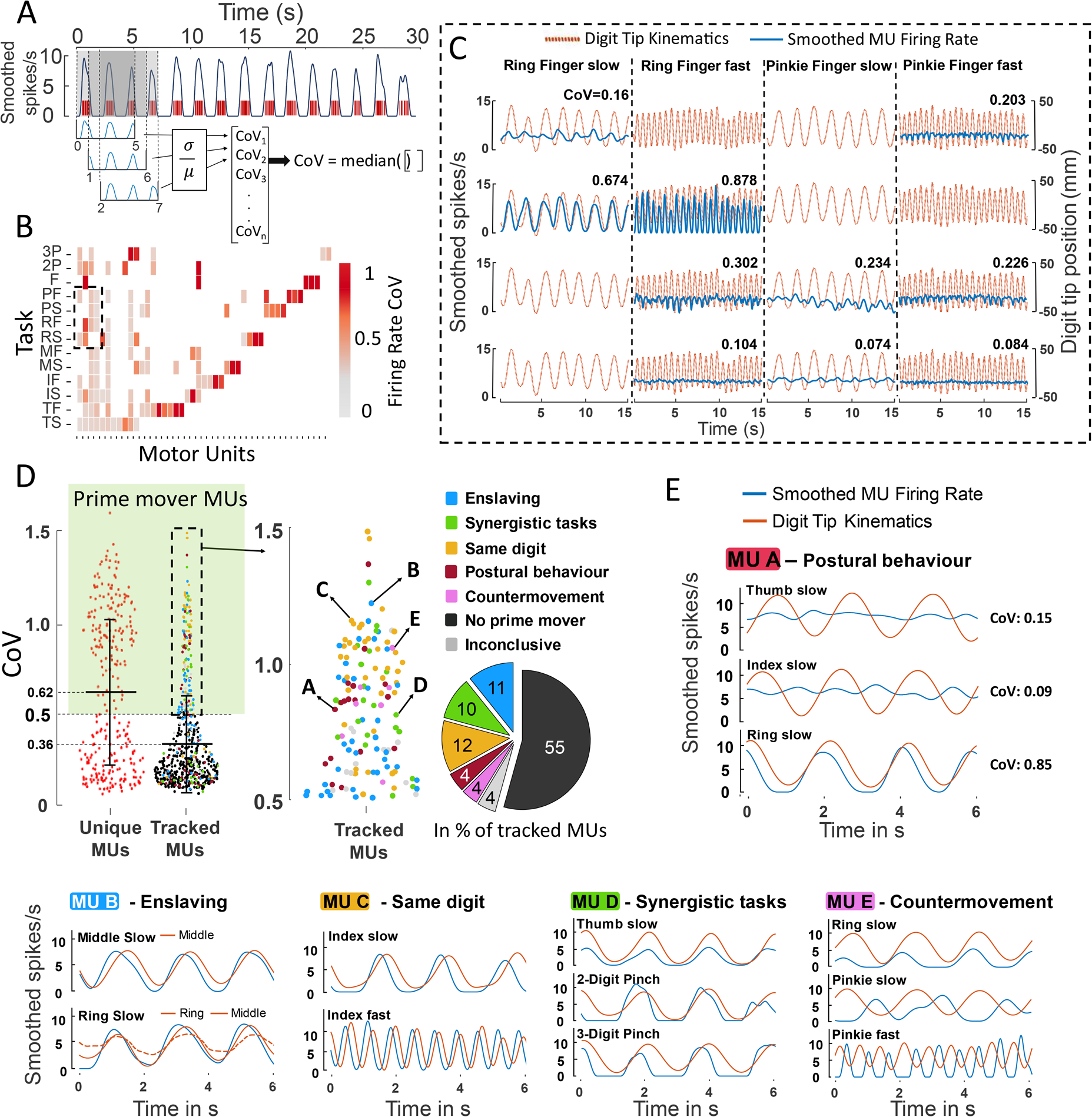
Prime mover firing behavior of MUs can be traced back to single-digit tasks of only one specific digit. (A) Coefficient of variation (CoV) as measure of task modulation of motor unit firing behavior. CoV is calculated as the median of all CoV values in moving windows of five seconds, in order to account for sections of the signal with inconsistently detected firings. (B) Matrix of all MUs of one exemplary participant. MUs in columns, tasks in rows. Fields in the matrix are colored according to the CoV of the firing behavior of the MU in the respective task. (C) Subset of (B) with visualization of smoothed MU firings (blue) and digit tip kinematics (red) for four tasks of four exemplary MUs with the respective CoV values. (D) Left: Distribution of CoV values of all MUs across all participants, separated into unique and tracked MUs. “Prime mover” MUs defined as MUs with a CoV value of >0.5. Unique MUs show higher average COV values compared to the tracked MUs. Right: Detail view of tracked MUs showing prime mover firing behavior. Close inspection of the firing behavior of every MU reveals strong task modulated firing activity in only one specific digit’s movement. (E) Exemplary cases of the kinematics and firing activity of tracked prime mover MUs in the identified tasks. <u>MU A:</u> Strong task modulation only present for one digit. <u>MU B:</u> Strong task modulation present in two tasks, but explainable by enslaving of one digit during another digit’s movement (here: middle digit moving in ring digit task). <u>MU C:</u> Strong task modulation in both tasks of same digit in the two different speeds. <u>MU D:</u> Strong task modulation in multiple tasks, however these tasks are mechanically synergistic, as the thumb is also part of the 2-digit and 3-digit pinch movements. <u>MU E:</u> Strong task modulation in multiple digits. Close inspection of the kinematics reveals opposite periods of MU activity with respect to flexion/extension phases (during extension for ring, but flexion for pinkie finger). This indicates a counter-activation of the one digit in the respective other digits task, possibly voluntarily by the participant in order to avoid enslaving.

The distribution of CoV values of all MUs across all participants revealed a higher average modulation of firing activity for MUs identified in only one task (unique MUs, mean CoV = 0.62 ± 0.42) compared to MUs tracked across multiple tasks (tracked MUs, mean CoV = 0.36 ± 0.31) (Fig. 3D). Unique MUs also showed a higher relative number of prime mover MUs compared to tracked MUs (52.1%, n= 147 vs. 24.5%, n=83) in single digit tasks. A detailed inspection of the firing patterns and kinematics revealed that tracked prime mover motor units only exhibited strong task modulated firing activity for one specific digit. The firing behavior of these MUs in other tasks they were identified in could be attributed to one or a combination of the following explanations (Fig. 3D-E):

#### Enslaving

The motor unit shows prime mover firing behavior for different digits A and B. Inspection of the corresponding digit tip kinematics reveals the presence of strong enslaving (mechanic coupling) of digit A, meaning digit A was following the movement of the actual digit B during digit B’s intended isolated motion (see Fig. 3E, MU B: Prime mover behavior in middle finger slow and ring finger slow tasks). The kinematics of middle fingertip during ring finger slow task shows strong middle finger movement (red dashed line). This phenomenon has been described previously and is attributed to musculoskeletal and neural constraints (Häger-Ross & Schieber, 2000; Kim et al., 2008; Santello et al., 2013; Zatsiorsky et al., 2000). We therefore assume the MU to be a prime mover MU only for digit A, with the strong task modulated activity during digit B’s movement attributed to the coupling of digits A and B.

#### Synergistic tasks

The motor unit shows prime mover firing behavior (CoV >0.5) for a single-digit task of digit A and one or multiple grasp tasks (see Fig 3E, MU D: Prime mover behavior for thumb slow task, as well as for 2-digit and 3-digit pinch tasks). As the grasps require movement of multiple digits in the same degree of freedom as for the respective single-digit tasks (synergistic movements), the prime mover firing behavior of the motor units in the can be attributed exclusively to the movement of digit A.

#### Same digit, different speed

The motor unit shows prime mover behavior in two single-digit tasks of the same digit A during both the fast and slow movement speed and show no prime mover behavior for other digits. The MU is therefore a prime mover MU for digit A (Fig 3E, MU C: Prime mover behavior for both index digit speeds).

#### Postural behavior

The MU was identified in single digit tasks of digits A, B, C and D, but shows prime mover behavior only for digit A and low task modulation for the remaining digits B, C & D (Fig. 3E, MU A: Prime mover behavior for thumb slow task, weak to no task modulation for ring digit slow & fast tasks). The MU is therefore a prime mover MU exclusively for digit A, with the weakly task-modulated firing activity during other digits movements attributed to digit A intentionally being kept in a stationary or postural position by the participant in order to prevent the enslaving of digit A during digits B, C and D’s movement.

#### Countermovement

Related to the postural behavior explanation above. MU shows prime mover behavior for digit A and medium to strong task modulated behavior for digit B (Fig. 3E, MU E: Prime mover activity for pinkie digit fast and slow tasks, also considerable task modulation for ring digit slow task). Closer inspection of the corresponding digit tip kinematics reveals the motor unit showing high activation during the opposite movement phase for digit B compared to digit A (In Fig. 3E, MU E: activation during flexion phase for pinkie digit, but during extension phase for ring digit. This behavior can be attributed to active voluntary prevention of enslaving by the participant, resulting in MU activation during the opposite phase to the active movement of digit B to keep digit A in a steady position.

When checking all 83 MUs that were tracked across multiple tasks and showed prime mover behavior in one or multiple of the movements for the application of the above conditions, 75 MUs could be assigned to one or a combination of multiple explanations (Fig. 3D). In these MUs, the prime mover firing behavior could be traced back to the movement of one specific single digit.

### Highly task-specific recruitment of prime mover motor units

We found only a small proportion of motor units showing prime mover activation (CoV >0.5) active in multiple tasks. Figure 4 A shows the ratio of tracked vs. unique motor units among different groups of movement tasks. A total of 97 ± 2.5% of motor units found in the first 20 seconds of a single digit task, were also found active in the second 20 seconds of the same task at the same movement speed. Looking at single digit tasks of the same digit, but at fast (1.5 Hz) and slow (0.5 Hz) movement speeds, only 17 ± 16.2% of prime mover MUs found at one speed, were also identified during the task of the respective other speed.

**Figure 4.**
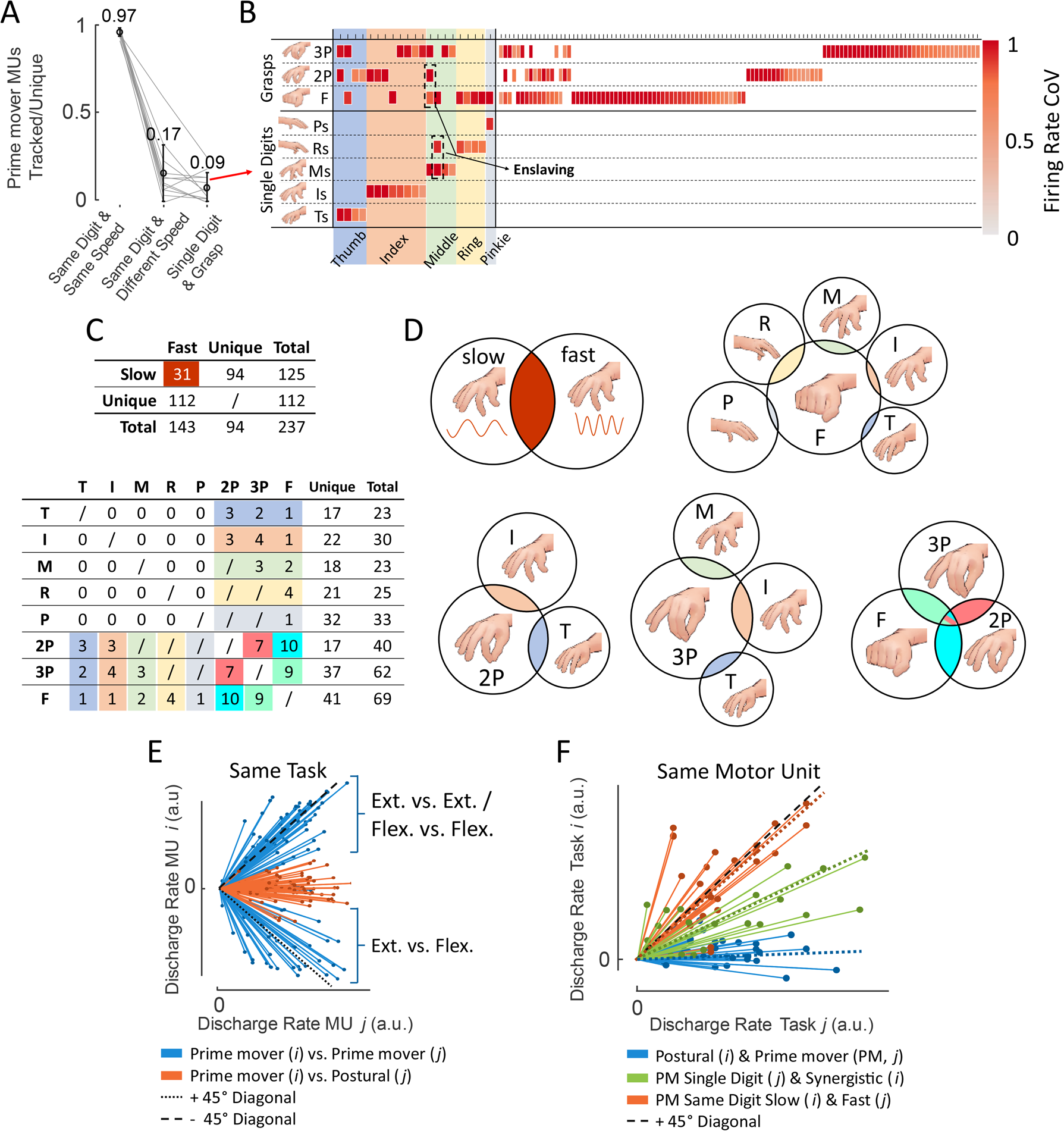
Highly task specific recruitment of MUs. (A) Ratio of unique vs tracked prime mover MUs within three groups of tasks: Identical single digit tasks (“Same Task & Same Speed”), Fast and slow speed of the same single digit n (“Same Task & Different Speed”) and within single digit tasks and synergistic grasping tasks (“Single Digit & Grasp”). Only a small proportion of motor units is shared between tasks of different speeds and single digit/synergistic tasks. (B) Matrix of all prime mover MUs identified in one or more of the three grasping tasks. Only a considerably small number of these MUs have also been identified during the single-digit tasks of the involved digits. (C) Top: Prime mover MUs identified in fast and slow movements of the same digit. Number of shared and unique MUs between speeds is shown. Bottom: Table of prime mover MUs identified in the single digit slow tasks and the synergistic grasp tasks. Number of shared MUs between grasps and involved single digits is shown. Note that there might be some hidden overlap between fields in cases where a MU is shared between more than 2 tasks (see Fig. 4B). (D) Prime mover MUs active during one digit speed or shared between both speeds, active only in grasp tasks and shared with the involved single digits, and unique/shared MUs within the grasp tasks. Area of the circles and circle overlaps are proportional to the number of motor units. Color coding is consistent within figures (B) – (D). (E) Linear regressions of scatter plots of MU discharge rates within a task reveal three main modules of neural input, namely digit flexion, digit extension and postural digit control. (F) Similar to (E), but linear regressions between discharge rates of the different tasks same motor units were identified in. Regressions split between cases where a unit was showing prime mover and postural behavior (blue), prime mover behavior in a single digit task and a synergistic grasp (green) and prime mover behavior during the two speeds of the same digit. Dotted lines represent the median regression line of each group.

Single digit tasks and synergistic grasping tasks involving the concerned digits shared an even lower ratio of prime mover motor units. Only 9 ± 8.2 % of MUs identified as prime movers during either a grasp task or single digit task were found active in both movements (Ratios obtained from averaging the single participant ratios). Since only a small proportion of MUs were found to be shared between movements of the same digits at different speeds, only the single digit movements performed at the same movement frequency as the grasp tasks (i.e., slow movements, 0.5 Hz) were considered. Figure 4 B illustrates these motor unit ensembles identified during the synergistic grasp tasks and respective single digit tasks. The color corresponds to the CoV values of the MUs in the concerned task (all >0.5). All prime mover motor units identified in the grasping tasks are displayed, together with all single digits tasks these MUs also served as prime movers. Enslaving of digits as described in the previous section can be observed in two cases. Figure 4 C shows the quantity of shared and unique prime mover motor unit ensembles during slow and fast single digit tasks (top), and within all combinations of slow single digit and grasp tasks (bottom). Only 31 of all 237 prime mover MUs found in the single digit tasks were active at both movement speeds.

Mechanically synergistic grasping tasks showed only minor overlap in MU ensembles with their involved single digits. Only 21 of all 283 prime mover MUs identified in the slow single digit tasks or grasp tasks were active during both movements. These values do not equal to the ratios shown in Figure 4 A, because Figure 4 A shows the average over all ratios obtained from the single participants to also show inter-participant variability, while the tables in Figure 4 C show the summation of all MUs found across all participants. It is important to note that in the synergistic grasp tasks, the digits moved in the same degree of freedom as during the single-digit tasks. In addition, we looked at the overlap between the grasp tasks. In total, 20 MUs were active as prime movers during two or all three grasp tasks, compared to 171 MUs identified in these tasks in total. Note that the numbers of shared MUs mentioned here do not equal exactly to the summation of fields of shared MUs in the table of Figure 4C, as there is some hidden redundancy in the numbers in cases where a MU was identified in more than two tasks, that cannot be displayed in a 2D table. All considered tasks showed a high number of MUs uniquely identified during each specific task. In Figure 4D, this low overlap in MU ensembles within certain groups of tasks is further visualized. The areas of the circles and overlaps are proportional to the number of MUs.

We find three main manifolds of neural input to the extrinsic hand muscles during the performed tasks: two manifolds controlling prime mover behavior, one for digit extension and one for digit flexion, both causing strong modulation of discharge activity with movement. The third manifold represents the postural control of digits, resulting in low movement-modulated firing behavior. Figure 4E shows this through the linear regressions of scatter plots obtained from plotting the normalized smoothed discharge rates of all motor units within the same task against each other, resulting in three clusters of regression lines. One shows clear positive slopes resulting from the scatter plots of two prime mover MUs encoding flexion or two prime mover MUs encoding extension. The second cluster is characterized by negative slopes obtained from scatter plots of two prime mover motor units encoding different movement periods (one encoding flexion, the other encoding extension). The last cluster shows flat slopes, generated from scatter plots of two motor units, where one is a prime mover MU, while the other showed postural firing behavior.

Figure 4F shows the same motor unit can be controlled by different manifolds. We plotted the linear regression lines of scatter plots similar to Figure 4E, but instead of the different motor units within the same task, we show the different tasks a motor unit was identified in. As already reported before, we observed that motor units can act both as prime mover motor units (CoV > 0.5) during one task, and postural motor units (CoV < 0.5) in other tasks (indicated in the plot by the blue group of regression lines, showing flat slopes). We also see that no motor unit can serve as a prime mover MU for both extension and flexion, as no regression line shows a clear negative slope. However, we found that prime mover motor units in a single digit task and a synergistic grasp task, usually exhibit higher discharge rates in the grasp tasks compared to the single digit tasks (Fig. 4F, green group of regression lines. Slope of median regression line clearly below the diagonal). The few motor units that were found active in both speeds of the same digit, showed similar discharge rates in both speeds (Fig. 3F, red group. Slope of median regression line close to the diagonal).

## Discussion

We recorded synchronized 3D hand kinematics and HD-sEMG data of 12 participants from 320 electrodes placed across the participant’s full forearms, recording muscle activation signals from most extrinsic hand muscles during 13 dynamic hand and digit tasks. Tasks included periodic single digit flexion and extension at two movement speeds per digit and three different grasps. Single motor unit activity was extracted from the HD-sEMG.

We found highly task-specific recruitment of prime mover MUs during single digit movements at different speeds and mechanically synergistic grasping tasks. Unexpectedly, across all participants only 17 ± 16.2% of prime mover MUs identified in a single digit task were active during both movement speeds. Therefore, distinct ensembles of motor units seem to be responsible for movements of same digits at different movement velocities.

Interestingly, an even smaller relative number of shared prime mover motor units was found between single digit tasks and mechanically synergistic grasping tasks. When considering only tasks performed at the same movement speed, only 9 ± 8.2 % of prime mover MUs were identified as shared between the grasps and corresponding single digit movements across all participants (Fig. 4A-D). On a motor unit level, dynamic multi-digit grasps and pinches are therefore more than the sum of their mechanical parts. Although the digits involved in the grasps moved in the same degree of freedom as during the single digit tasks, the small overlap in motor unit ensembles could be attributed to the higher level of precision required in multi-digit grasps in these unconstrained dynamic conditions, as accurate coordination especially of the thumb and index finger is required for fingertip-to-fingertip contact. To facilitate such fine movements, the neural system seems to recruit additional, previously unrecruited motor neurons compared to less precise single digit flexion/extension movements, where no specific point in space must be reached by the digit.

During single-digit movements, we found strong evidence supporting the hypothesis that distinct movement-encoding prime mover motor units control each digit. All occurrences of prime mover behavior of one motor unit in multiple single-digit tasks could be attributed to one or a combination of multiple explanations, like the synchronous movement of multiple digits due to mechanical coupling (enslaving) or active prevention of such by the participant (Fig. 3D,E). It should be noted that the participants were not explicitly instructed to prevent or allow such synchronous movements of multiple digits during single-digit tasks. Zatsiorsky et al., 2000 reported digits experiencing enslaving to produce forces of up to 67.5% of their isolated maximum forces during isometric contractions and it is suggested to be rooted in neural interactions in the underlying neurophysiological system (Häger-Ross & Schieber, 2000; Kim et al., 2008; Santello et al., 2013; Zatsiorsky et al., 2000).

The vast majority of motor units that could be tracked across multiple digits showed strong task modulation only for a single digit, with low modulation of firing activity in the remaining tasks. Only 4% of tracked motor units showed firing activity across tasks that could not be traced back to the prime mover behavior of a single digit. This could be attributed to false positive matches during the tracking of MUs across tasks, although we chose a decision boundary of cross-correlation values specifically aimed to prevent such false positive matches. Additional explanations outside of the ones proposed in figure 3D and 3E could be present as well.

Looking at MUs shared between the three grasp tasks, we found a range of 7.4% (2P&3P and 3P&F) to 10.0% (2P&F) of shared prime mover MUs. Tanzarella et al., 2021 found percentages of shared motor units between grasping tasks of <25% in isometric conditions. This difference in observed share can again be attributed to the dynamic dimension of the present dataset, that introduces a complexity to the performed movements that is not present in isometric conditions and might lead to a higher task-specificity of recruited motor unit ensembles.

Overall, all observed movement tasks were driven by a high share of entirely task specific prime mover motor units. This fits recent findings where a convolutional neural network was trained to continuously predict the 3D kinematics of all hand and digit joints from the HD-sEMG data of the same dataset used in this study. Analysis of the high dimensional latent space vectors of the neural network into 2D and 3D reveals distinct clusters of single digit and grasp tasks, with clearly separated clusters between the fast and slow movement speeds (Sîmpetru et al., 2022). It needs to be considered that in all results, high inter-participant variability could be observed, that should be investigated further.

As the decomposition method used here is validated and generally used in isometric movements, special care was taken to include only valid motor units in the analysis. We tracked each participant’s motor units across tasks based on the cross correlation R of their motor unit action potential shapes. We used groups of known identical MUs (true positives) and known distinct MUs (true negatives) to determine a decision boundary based on their R values. The clear clustering of cross-correlation values of the true positive and true negative groups shown in Figure 2A confirms the validity of the tracking approach. The clustering was even more pronounced as observed by Tanzarella et al., 2021, where a similar approach for validating the tracking of motor units across tasks was performed. This can be explained by the higher pulse-to-noise ratio threshold applied here for the inclusion of motor units in the tracking approach (30 dB vs 25 dB in Tanzarella et al., 2021, average 34.9 ± 1.6 dB vs 30.7 ± 4.6 dB). During single-digit movements, we found strong evidence supporting the hypothesis that distinct movement-encoding motor units (prime mover MUs, Coefficient of variation of smoothed firings > 0.5) control each digit. We showed that a MU acts as a prime mover for not more than one single digit.

The presented results can have a strong impact on the field of neurorehabilitation, in particular the control of exoskeletons and prostheses. As shown recently in a single participant (Ting et al., 2021) and a larger cohort of patients with real-time motor unit feedback (Oliveira et al., 2022), motor units below the level of the lesion of motor-complete spinal cord injury patients still receive neural input and can be decoded by non-invasive neural interfaces. Distinct pools of motor neurons encoding digit movements, when decoded in real-time, can allow for more precise and intuitive control of assistive devices in the future. Further research building on the presented findings could investigate the neural control of motor unit pools during synergistic hand movements even further with including a larger variety of behavioral tasks and with special focus on participant-specific variability to explore the generalizability of the results. A closer look at the motor units in terms of recruitment thresholds and firing rate in single digit movements and mechanically synergistic movements could yield valuable insight in possible flexible control of motor units during dynamic hand movements.

All above conclusions come with the limitation that with the current methods of motor unit decomposition from surface and current intramuscular EMG signals, only a part of all active motor units can be identified. However, as this limitation applies to all movements performed in this study in the same way, we can assume the qualitative validity of the above discussed results. With a higher number of identified motor units, through high-density intramuscular EMG (Muceli et al., 2015, 2022) or improvements in decomposition methods of surface EMG signals, the observed shared pool of motor units between single digit movements and grasping tasks, as well as between different grasping tasks, might increase.

To sum up, we decoded motor unit activity during single- and multi-digit hand movements with synchronized 3D hand kinematics. Motor units either showed prime mover firing activity, showing strong modulation in discharge rate with movement or were considered as postural motor units. Depending on the current movement task, the same motor units can either act as prime mover or postural MUs (Fig. 5). We found the movement of single digits to be controlled by distinct pools of prime mover motor units that are not shared between digits. Unexpectedly, most prime mover MUs were not active during both speeds of single digit movements. This highly task specific recruitment of motor units also extended to multi-digit pinch and grasping movements. We observed only minor overlap in recruited prime mover motor units of the grasps with their involved single digits (Fig. 5). Supplementary video S2 shows these findings in exemplary tasks of two subjects with the corresponding synchronized experimental videos and 3D kinematics. These results have significant implications on the field of motor control of synergistic hand movements, as well as the development of neural interfaces for the control of prosthetics and assistive devices.

**Figure 5.**
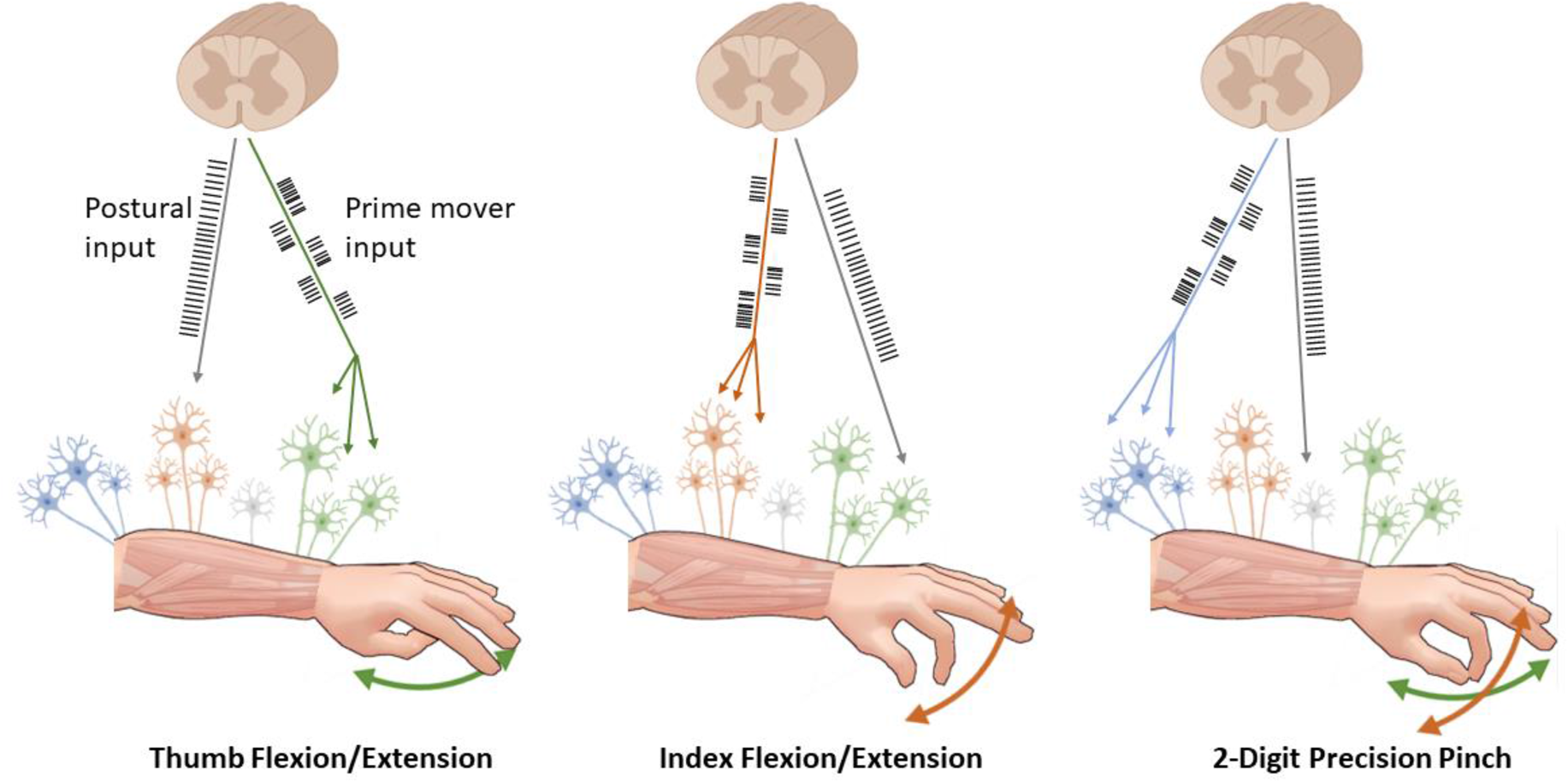
Summary of results. We observe highly task-specific recruitment of movement encoding motor units. Single digit movements are controlled by distinct groups of motor units. Motor units are either showing prime mover behaviour associated with strong movement-correlated activity, or postural activation. The same motor unit can show both behaviours, depending on the present movement task. Mechanically synergistic movements do not share prime mover motor units with the single digit movements involved in the synergistic task.

## Materials and Methods

### Experimental Setup

Twelve healthy participants (age= 25.9 ± 2.8, ten males, two females) volunteered in this study. The consent form and study protocol were approved by the Friedrich-Alexander University Erlangen-Nürnberg ethics committee (n. 150 21 B) in accordance with the Declaration of Helsinki. HD-sEMG data was recorded from five 64-channel electrode grids (three 8×8 grids, interelectrode distance (IED) 10 mm and two 13×5 grids, IED 8 mm. OT Bioelettronica, Turin, Italy). The electrode grids were placed on the participant’s proximal and distal forearm to record data from all extrinsic hand muscles, particularly the m. flexor digitorum superficialis and profundus, m. extensor digitorum, m. extensor pollicis longus and brevis and m. flexor pollicis longus (see Fig. 2B). Monopolar EMG signals were recorded at a sampling frequency of 2048 Hz and filtered with a 10-500 Hz bandpass filter by the acquisition system (Quattrocento, OT Bioelettronica, Turin, Italy). During the experiments, the participants stood upright with their elbow flexed at a 90° angle resulting in their forearm and digits pointing forward, with the palm perpendicular to the ground and neutral wrist position (see Fig. 1A). All participants used their dominant hand. A realistic virtual hand model embedded with 23 anatomical degrees of freedom was developed (Blender 3.0, Blender Foundation). The virtual hand can be configured to display realistic single or combined hand and digit movements at precise predefined movement speeds. During the experiment sessions, this hand was displayed on a monitor in the center of the participant’s field of view (Fig. 1A). The participants were instructed to imitate the shown movement as precisely as possible regarding the range of motion and movement speed to ensure comparable execution of movements across tasks and participants. Before each recording, the participants could practice the tasks to get familiarized with the movement and movement frequency.

Each participant performed the same 13 hand and digit tasks for 45 seconds per task. 10 tasks consisted of periodic single-digit flexion and extension of the metacarpophalangeal (MCP) joint. For each digit, two separate tasks were performed at frequencies of 0.5 Hz (slow) and 1,5 Hz (fast). 1 Hz corresponds to one cycle of full extension to full flexion to full extension per second. In addition to the single-digit tasks, three different grasp tasks were performed, consisting of a five-digit power grasp, a two-digit precision pinch using the thumb and index finger, and a three-digit precision pinch utilizing the thumb, index finger and middle finger (Fig. 1D). Like the single-digit tasks, the grasps were performed periodically at a frequency of 0.5 Hz or 2 seconds from full grasp to default position to full grasp.

### Markerless 3D Kinematics Acquisition

During the experimental session, the participant’s hand was placed in the center of the measurement volume of a camera setup, recording the hand and digit movements with five industrial RGB cameras from different angles and positions (Cameras: DFK-37BUX287, Objectives: TCL 0814 5MP; The Imaging Source, Charlotte, USA). A microcontroller (Arduino UNO R3) was used to send an analogue trigger signal at a frequency of 140 Hz to all cameras for precisely synchronized frame acquisition at a framerate of 140 frames per second. The trigger signal was also sent to the HD-sEMG acquisition system for subsequent synchronization of video and EMG data.

For obtaining the 3D kinematics of the hand and digit movements performed by the participants, the open-source Python toolboxes *DeepLabCut* (DLC) and *Anipose* were used. DLC offers a framework for markerless and automatic pose estimation of sets of videos through custom training of pre-trained neural networks (Mathis et al., 2018; Nath et al., 2019) and has been used previously for motion tracking in neuroscience research (Del Vecchio, Sylos-Labini, et al., 2020; Kane et al., 2020; Sehara et al., 2021; Solby et al., 2021) and validated against state-of-the-art motion tracking systems (Kosourikhina et al., 2022; Van Hooren et al., 2023). The custom neural network used for this study was trained on a manually labeled dataset of a wide variety of hand postures generated prior to the experiment on the base of a pre-trained ResNet-101. Figure 1A shows the labeling scheme chosen for automatic posture tracking in this study. 21 labels located on the five digit tips, the distal interphalangeal (DIP), proximal interphalangeal (PIP), and metacarpophalangeal joints (MCP) of the index to pinkie finger, the interphalangeal (IP) and MCP joints of the thumb and the radiocarpal wrist joint, allow for full representation of the physiological number of degrees of freedom of hand and digit movements. The trained neural network was then used to track the 2D kinematics of the hand and digits in the videos of all five cameras. With *Anipose* we then triangulated the 2D label positions into the 3D hand kinematics. The toolkit provides functions for calibrating the extrinsic and intrinsic parameters of the camera setup and temporal and spatial filters for fine-tuning the triangulation process (Karashchuk et al., 2021).

For an evaluation of the accuracy of the automatically tracked and triangulated 3D hand kinematics, the 2D images of 100 representative hand postures taken from 10 participants were labeled first manually by two experts and then automatically by the same previously trained deep neural network used for the tracking of the experiment videos. Both datasets were triangulated using the same camera calibration and filter parameters, and the 3D Euclidean distance between manually and automatically tracked posture was calculated. The averaged 3D position of the postures obtained from the two experts’ manually labeled images was considered the ground truth data. The 3D label positions derived from the markerless tracking by the neural network showed a mean absolute error (MAE) of 2.90 ± 2.07 mm to the ground truth positions, with 3.27 ± 2.23 mm MAE to scorer 1 and 3.38 ± 2.74 mm MAE to scorer 2, respectively (Fig. 1B). The cumulative distribution function (CDF) of the deviation of the automatically tracked 3D position to the ground truth was computed as an estimate of the probability of any label deviating by a specific distance from the correct 3D position (Fig. 1B).

### Motor Unit Analysis

All offline analysis and EMG signal processing were performed using MATLAB (MATLAB R2022b, MathWorks Inc., Natick, MA). After data acquisition, the EMG data was cleaned from channels showing outlier behavior and poor signal-to-noise ratio in a semi-automatic way. Channels with exceptionally bad signal-to-noise ratios were removed automatically, followed by a careful visual inspection of the signals to remove additional channels. After the automatic and manual inspection, the average relative number of deleted EMG channels per participant was 11% of the full 320 channels. The EMG data was cleared from the 50 Hz power line interference noise and its 100 Hz and 150 Hz harmonics with a notch filter.

To extract the discharge timings of the individual motor units, HD-sEMG data were decomposed using an approach based on blind source separation and convolutional kernel compensation that has been described and validated before (Del Vecchio, Negro, et al., 2019; Farina & Holobar, 2016; Holobar & Zazula, 2007; Negro et al., 2016). To assess the accuracy of MU identification, we calculated the pulse-to-noise ratio (PNR) as in (Holobar et al., 2014). Only motor units with a PNR of >30dB were considered for further analysis. This PNR has been reported as a suitable threshold in the longitudinal identification of motor units (Vecchio & Farina, 2019). Experts reviewed all motor unit pulse trains manually and corrected all false-negative and false-positive detected firings as described by Del Vecchio, Holobar, et al., 2020. Each motor unit’s action potential (MUAP) shape was extracted by spike-triggered averaging (Del Vecchio et al., 2017; Farina et al., 2002) for each channel of the HD-sEMG grids. To ensure stable MUAP shapes across the entire signal duration, we excluded all MUs with <100 action potentials detected in any given task.

For additional validation of the estimation of the discharge timings in the decomposition process, the MUAP waveforms and consistency of MU location were visually evaluated throughout the signal via shimmer plots (Fig. S1). Centered at each spike instant with a window of 25ms, HD-sEMG signals were extracted to visually inspect whether individual MUAP shapes for each instance in the spike train agree with the template waveform of the MU. This method assumes that the unique MU waveform is largely preserved throughout the contraction. Additionally, we looked at the location of each MU within the HD-sEMG grids to ensure distinct motor unit territories (Fig. S1).

The spike trains of each MU for each task were smoothed with a Hann-window as proposed in Negro et al. (2009) with window length1s for fast tasks (movement frequency 1.5 Hz) and 0.33 s for slow tasks (0.5 Hz). Different window lengths were chosen to account for the difference in flexion/extension frequency of the tasks and should ensure comparable CoV values between fast and slow movements. As a measure of task modulation of MU firing behavior for each individual task, the coefficient of variation (CoV, defined as the signals standard deviation σ / signals mean μ) of the smoothed motor unit discharges was computed. As some MUs contained short periods of undetected firings due to actual physiological inactivity or firings not detected by the decomposition method, the CoV was computed as the median value of all CoV values obtained in windows of 10s (fast tasks) or 3.3s (slow tasks), shifted by 1s between windows (Fig. 3A).

### Motor Unit Tracking

A 2-D inverse rectangle (IR) spatial filter was applied to the MUAP data for noise reduction, especially important in low-amplitude MUs (Farina et al., 2003). The identification of motor units across movement tasks makes use of the consistent placement of the electrode grids with respect to the muscle’s position, as grids were not removed in between tasks. We also assume the change of volume conductor properties of the muscles did not change dramatically during the hand and digit movements, which would result in substantially altered MUAP shapes during and between tasks. Therefore, MUAP shapes of one motor unit in two separate tasks can be considered nearly identical in shape and similar in amplitude. As previous studies tracking MUs longitudinally used data acquired in isometric conditions and from a substantially lower number of HD-sEMG channels, we modified the previously proposed method of cross-correlation of the full grid of MUAP shapes with correlation values of 0.7-0.8 as a threshold for identical motor units (Del Vecchio, Casolo, et al., 2019; Martinez-Valdes et al., 2017; Vecchio & Farina, 2019). To determine the similarity of MUAP shapes of a pair of MUs, we first determined the 10 channels with the highest peak-to-peak MUAP amplitudes for each MU (MU A and MU B). We then computed the cross-correlation of MUAP shapes only between the 10 highest peak-to-peak channels of MU A with the same channels of MU B and vice versa.

The mean of these values was then determined as the final cross-correlation value R for each pair of MUs. This method considers not only the similarity of MUAP shapes, but also the motor unit specific territory that is expected to be consistent across tasks for identical motor units. To obtain a reliable R-value threshold for a positive MU match, we computed the distribution of R values within groups of known identical MUs (true positives) and known distinct MUs (true negatives). For the true positive values, all MUs in the dataset were matched with themselves by computing the MUAP shapes within two distinct 20s periods of the signals and performing the cross-correlation between both periods. As the decomposition result of each task was carefully checked manually by experts for identical MUs, the MUs within the tasks were considered as true negative matches. As shown in Figure 2A, the distribution of cross-correlation R values between the true positive and true negative groups shows clear clustering (true positive: R = 0.928 ± 0.042, true negative: R = 0.475 ± 0.212). To ensure no false positives during the matching of the MUs between tasks, the threshold value R_T_ was determined as R_T_=0.90, which excluded 100% of true negatives and included 80.4% of true positive matches. As a further validation, we computed the absolute difference in peak to peak amplitudes (ΔP2P) of each pair of MUs of the full dataset and the true positive and true negative groups. We observed clear clustering of (ΔP2P) values between the groups.

## Supplemental Material

**Figure S1.**
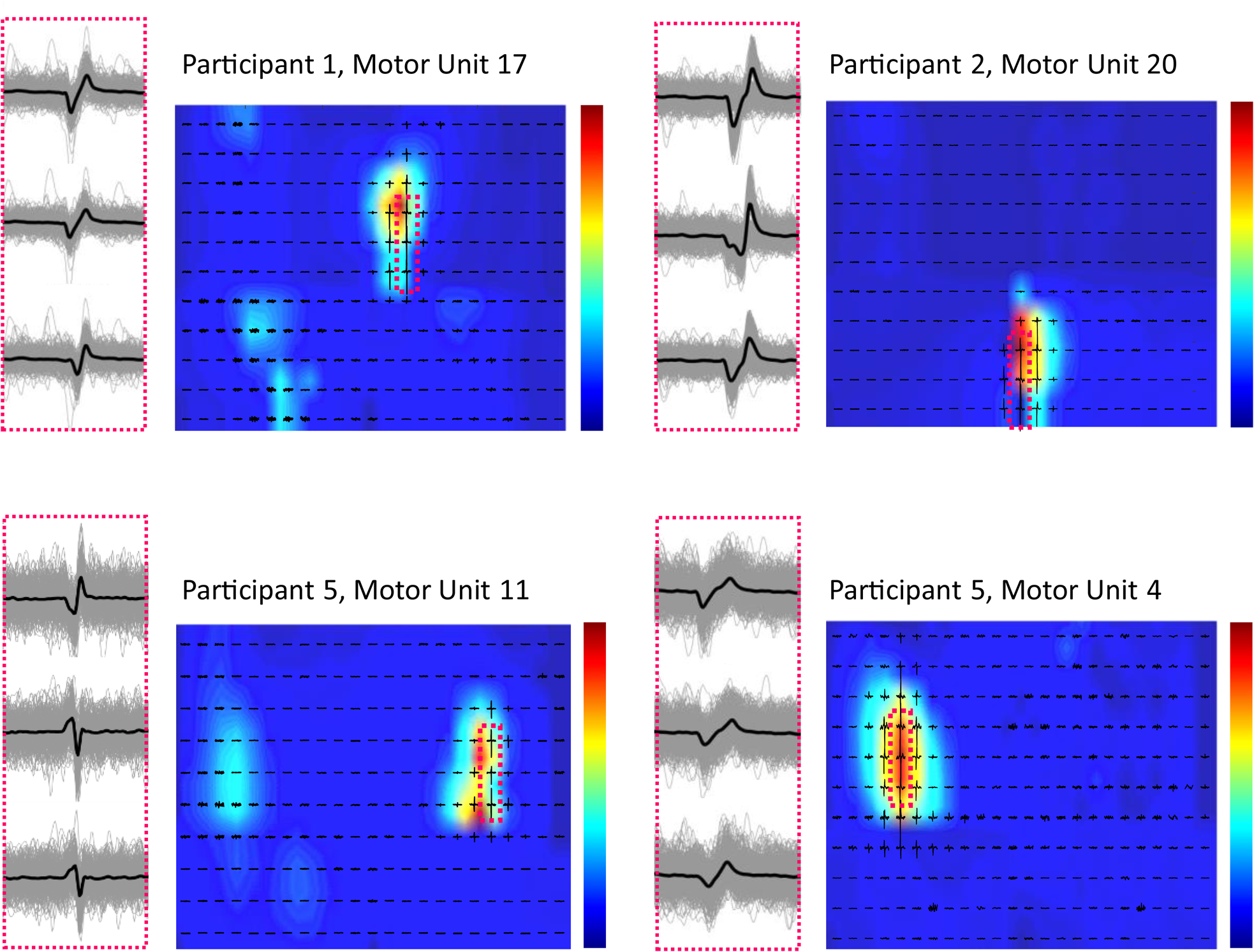
Evaluation of decomposition accuracy. Spike triggered averaging and heatmaps of MU activation within the HD-sEMG grids for four exemplary MUs in the full dataset. Left: Each individual spike of MU spike train in each channel of the HD-sEMG data superimposed, with black line representing the average motor unit action potential. Right: Spatial representation of motor unit territory. Colors represent the root-mean-square of the motor unit action potential across the full duration of the signal. Distinct centers of activation throughout the signal can be observed for all MUs.

### Video Link

**Video S2.** Video showing experimental videos and tracking of 3D Kinematics with synchronised single motor unit discharges of exemplary tasks and participants. Motor unit data shows task specific recruitment of prime mover motor units within single digit tasks, and between single digit and mechanically synergistic grasping tasks. Motor units can be acting as prime mover or postural MUs, depending on present movement task.

