## Supplementary figures and images for "Mechanical Hand Synergies during Dynamic Hand Movements are Mostly Controlled in a Non-Synergistic Way by Spinal Motor Neurons"

### Supplemental Figure 1

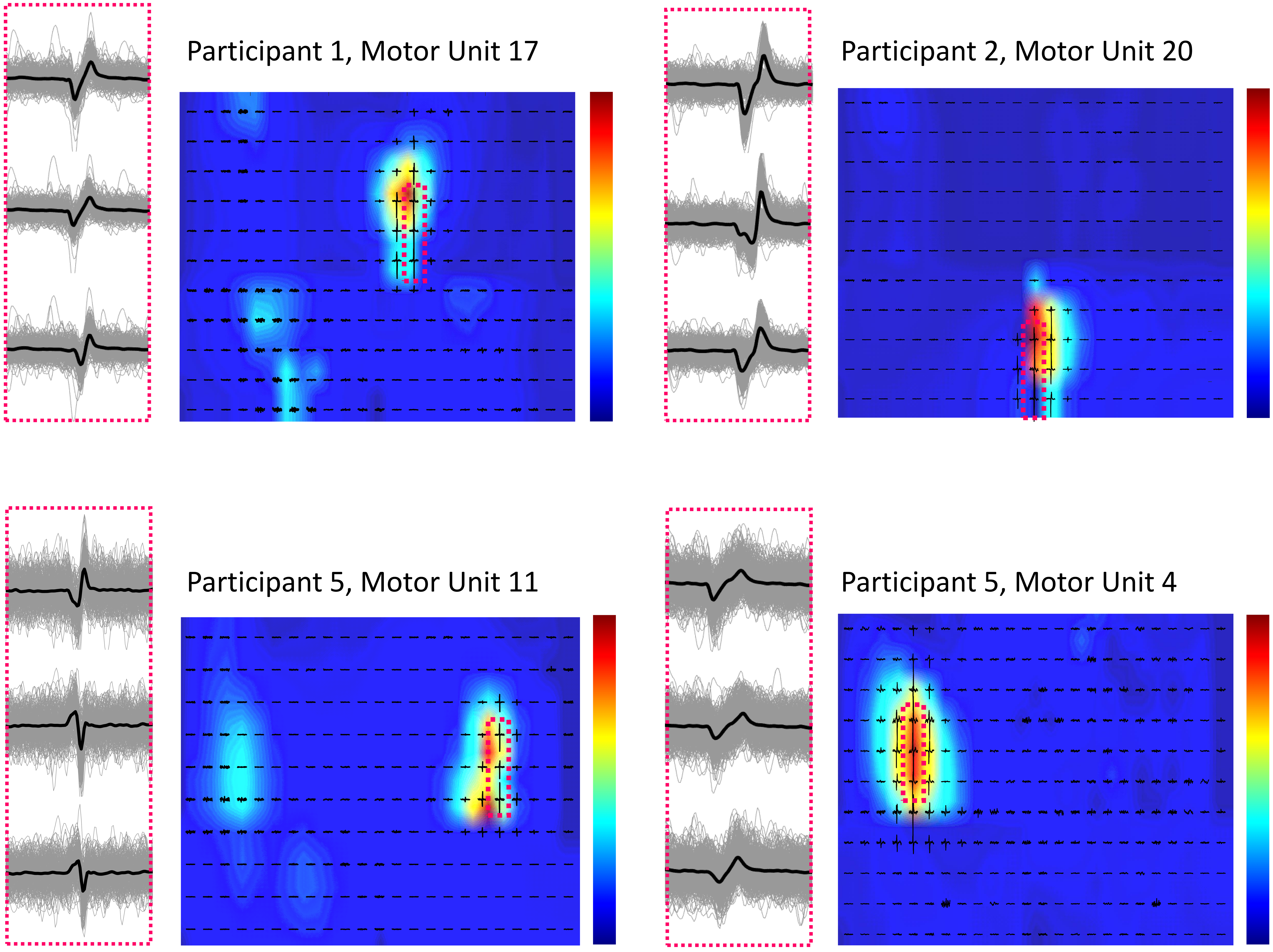
